# A multi-organ metabolic model of tomato predicts plant responses to nutritional and genetic perturbations

**DOI:** 10.1101/2021.09.30.462630

**Authors:** Léo Gerlin, Ludovic Cottret, Antoine Escourrou, Stéphane Genin, Caroline Baroukh

## Abstract

Predicting and understanding plant responses to perturbations requires integrating the interactions between nutritional sources, genes, cell metabolism and physiology in the same model. This can be achieved using metabolic modeling calibrated by experimental data. In this study, we developed a multi-organ metabolic model of a tomato plant during vegetative growth, named VYTOP (Virtual Young TOmato Plant) that combines genome-scale metabolic models of leaf, stem and root and integrates experimental data acquired from metabolomics and high-throughput phenotyping of tomato plants. It is composed of 6689 reactions and 6326 metabolites. We validated VYTOP predictions on five independent use cases. The model correctly predicted that glutamine is the main organic nutrient of xylem sap. The model estimated quantitatively how stem photosynthetic contribution impact exchanges between the different organs. The model was also able to predict how nitrogen limitation affects the plant vegetative growth, and to predict the metabolic behavior of transgenic tomato lines with altered expressions of core metabolic enzymes. The integration of different components such as a metabolic model, physiological constraints and experimental data generates a powerful predictive tool to study plant behavior, which will be useful for several other applications such as plant metabolic engineering or plant nutrition.

**One sentence summary:** A multi-organ metabolic model of tomato gives biological insights into the functioning of a plant such as xylem composition, the role of the stem and the response to environmental or genetic perturbation.

## Introduction

Systems biology can predict and explain how a whole organism responds to a stimulus or a perturbation and how different components of the organism are connected. In particular, one way to unravel physiological mechanisms is the use of constraint-based metabolic modeling. Constraint-based metabolic modeling relies on genome-scale metabolic networks, which gather all the identified metabolic reactions of an organism according to its genomic sequences and current metabolic knowledge summarized in databases (Gu, Kim, Kim, Kim, & Lee, 2019). It predicts and quantifies the metabolic pathways used thanks to mathematical constraints that encompass physiological states, such as the uptake rate of nutrients present in the environment. Initially developed for unicellular organisms, constraint-based metabolic modeling has been extended to diverse plant species such as *Arabidopsis thaliana* (Gomes de Oliveira Dal’Molin, Quek, Saa, & Nielsen, 2015; Mintz-Oron et al., 2012; Shaw & Cheung, 2018), *Medicago truncatula* (Pfau et al., 2018), barley (Grafahrend-Belau et al., 2013), *Steraria viridis* (Shaw & Cheung, 2019) and soybean (Moreira et al., 2019). These studies analyzed mechanisms such as the effect of the nitrogen source (Arnold & Nikoloski, 2014; Arnold, Sajitz-Hermstein, & Nikoloski, 2015; Gomes de Oliveira Dal’Molin et al., 2015; Seaver et al., 2018), the nitrogen fixation by symbiotic bacteria (Pfau et al., 2018), the effect of diurnal cycles (Gomes de Oliveira Dal’Molin et al., 2015; Shaw & Cheung, 2018) or the genetic modifications to perform in order to enhance a product of interest such as vitamin E (Mintz-Oron et al., 2012; Saha, Suthers, & Maranas, 2011). Some of these studies were supported with labeling data, such as Robaina-Estévez et al. (2017), which studied the metabolic differences between guard cells and mesophyll cells. For a full review of all models of plants developed so far, the reader is referred to (Gerlin, Frainay, Jourdan, Baroukh, & Prigent, 2021; Shaw & Cheung, 2020).

The representation of the multi-organ structure of plants through modeling is challenging (Clark, Guo, Morgan, & Schwender, 2020), but is required to better understand plant metabolism. To model exchanges between organs, a global exchange pool is usually used (Gomes de Oliveira Dal’Molin et al., 2015; Shaw & Cheung, 2018), but the veracity of the exchanged fluxes is not studied. Another major limitation is the lack of experimental calibrations since most of the methodologies gather heterogeneous data from literature, issued from multiple species and different growth scenarios, which could imply bias and inaccuracies. In addition, tomato (*Solanum lycopersicum*), a major agricultural crop, did not benefit of the advances in multi-organ metabolic models, as only a genome-scale metabolic model of leaf cell was published so far (Yuan, Cheung, Poolman, Hilbers, & van Riel, 2016), as well as a metabolic model of the fruit (Colombié et al., 2017, 2015; Li et al., 2020).

In this study, we aimed to develop a multi-organ metabolic model of tomato plant during vegetative growth, in order to provide to the plant research community a comprehensive tool adapted for many biological questions such as the impact of environmental factors on the early development of the tomato plant, the exchange fluxes of matter between organs, or the prediction of the reactions to be modified to obtain a desired phenotype. This model, named VYTOP (Virtual Young TOmato Plant), includes leaf, stem, root and, contrary to existing multi-organ models, dissociated xylem and phloem compartments. It is able to simulate all the major metabolic reactions used in the plant and the sink/source relationships between organs. The model was calibrated with homogenous data from experiments we performed, using an automated phenotyping platform, gathering both physiological data (growth, transpiration) and metabolomics (xylem sap chemistry, organ biomass composition). We chose five independent use cases to validate the usefulness of our model for a broad range of biological questions: i) prediction of core metabolic fluxes in each tomato organ, ii) contribution of stem to photosynthesis, iii) xylem composition, iv) impact of nitrogen limitation on growth, v) predictions of metabolic changes in transgenic lines. Beyond its use to model for the first time the fluxes between the organs of the tomato, this model could serve as a template for modelling other plants of interest as well as predicting and understanding the impact of diverse perturbations.

## Results and Discussion

### VYTOP, a multi-organ metabolic model

The multi-organ metabolic model, named VYTOP (Virtual Young TOmato Plant) (**Figure *1***) was built by aggregating genome-scale metabolic models of each organ. The published tomato leaf genome-scale metabolic model iHY3410 (Yuan et al., 2016) was used as starting point. We curated the model to integrate missing pathways such as the catabolism of different metabolites (lysine, ethanol, isoleucine, beta-alanine, leucine, cysteine). As the iHY3410 metabolic model IDs mostly followed the MetaCyc IDs nomenclature (Caspi et al., 2016), we converted them into the BiGG format (King et al., 2016) using a semi-automatic conversion framework. BiGG is widely used for genome-scale metabolic models from bacteria to microalgae and humans (Norsigian et al., 2020). The converted model was also manually curated to achieve mass balance for carbon, nitrogen, oxygen, phosphate and sulfur for each reaction. We name this novel tomato metabolic model Sl2183; it includes 2183 reactions, 2097 metabolites and 3433 genes. It is available in different formats: table format (Supplementary File 1), SBML format (Supplementary File 2), in the database BioModels (Glont et al., 2018) under the ID XXX [once the paper is accepted], and in the MetExplore database to enable pathway visualization and omics mapping (Cottret et al., 2018) https://metexplore.toulouse.inrae.fr/metexplore2/?idBioSource=6237.

To transpose this genome-scale metabolic model into a multi-organ model, we considered the whole plant as three main organs (Figure 1). Each organ was modeled with a replicate of the Sl2183 genome-scale metabolic model with an organ-specific biomass equation. In addition, each organ has specific physiological roles: photosynthesis/organic matter production in leaf, partial photosynthesis in stem and minerals/water uptake in root represented by accurate physiological constraints. The transport tissues xylem and phloem were defined as exchange compartments: xylem represents exchanges from root to leaf, phloem exchanges from leaf to root. Stem can exchange with both compartments in the two directions: uptake or contribution to xylem and phloem. Leaf and stem assimilate photons (i.e light) while root assimilates water and minerals. An ATP cost was set up for transport reactions between organs and xylem/phloem to include the transfer costs. The final VYTOP model includes 6689 reactions and 6326 metabolites.

**Figure 1:**
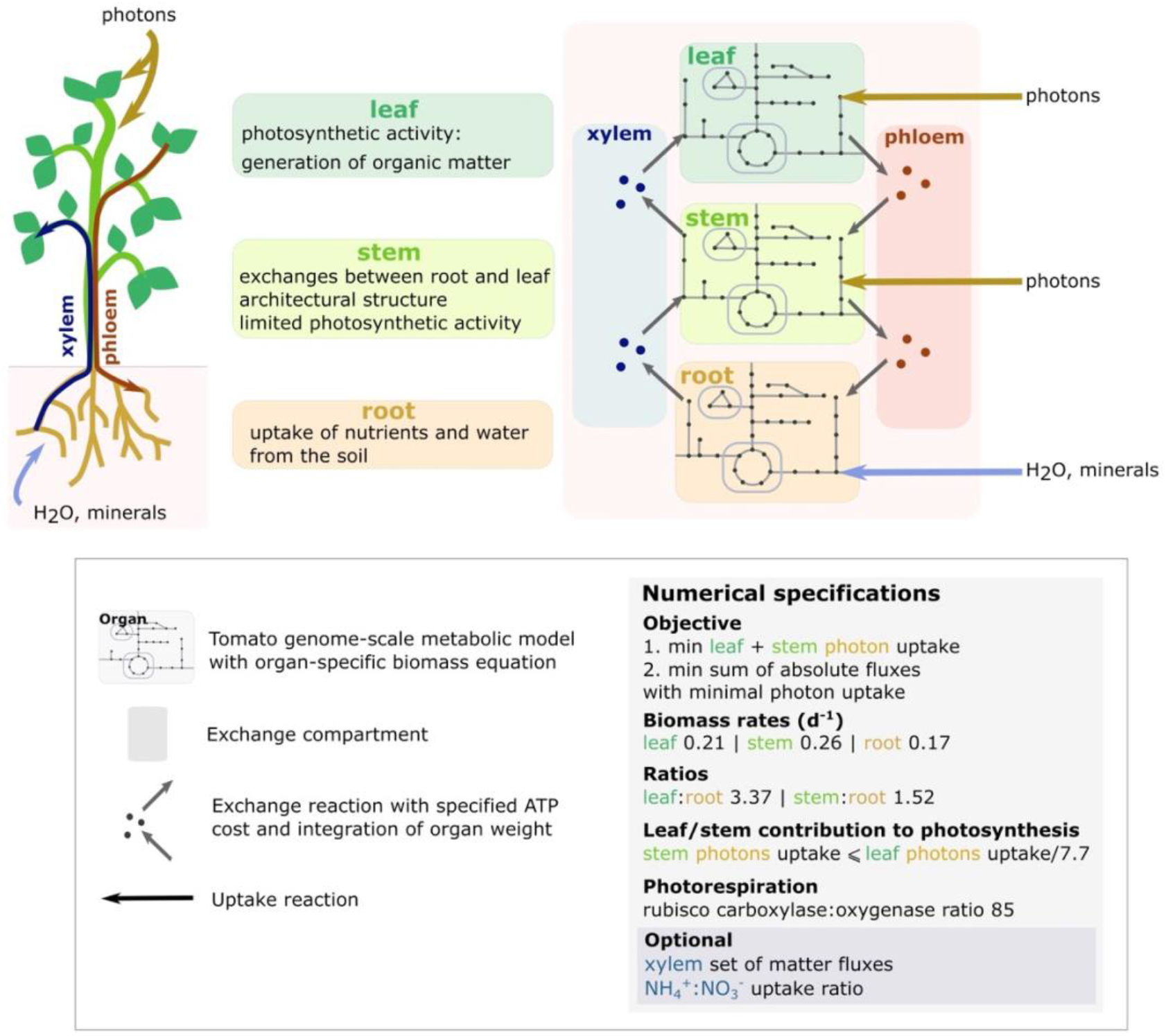
Generation and calibration of VYTOP (Virtual Young TOmato Plant) based on a tomato genome scale metabolic model. Top left panel: role of each organ as modelled in VYTOP. Top right panel: Schematic view of the different organs and exchange compartments as modelled in VYTOP. Bottom panel: legend, numerical constraints and objective functions used in VYTOP.

### VYTOP calibration with physiological and metabolic data

To calibrate VYTOP, we performed physiological and biochemical measurements (Figure 2) on 90 tomato plants. The growth of 90 plants was monitored on an automatic phenotyping facility, which allowed an automatic monitoring of plant transpiration rate. Sampling of 6 plants per day was performed during 10 days, starting at 28 days after seedling. Xylem metabolite concentrations and organ weights were determined on sampled plants. The biomass composition of each organ was measured in another experiment performed in the same environmental conditions, with the same tomato variety and a similar plant age.

**Figure 2:**
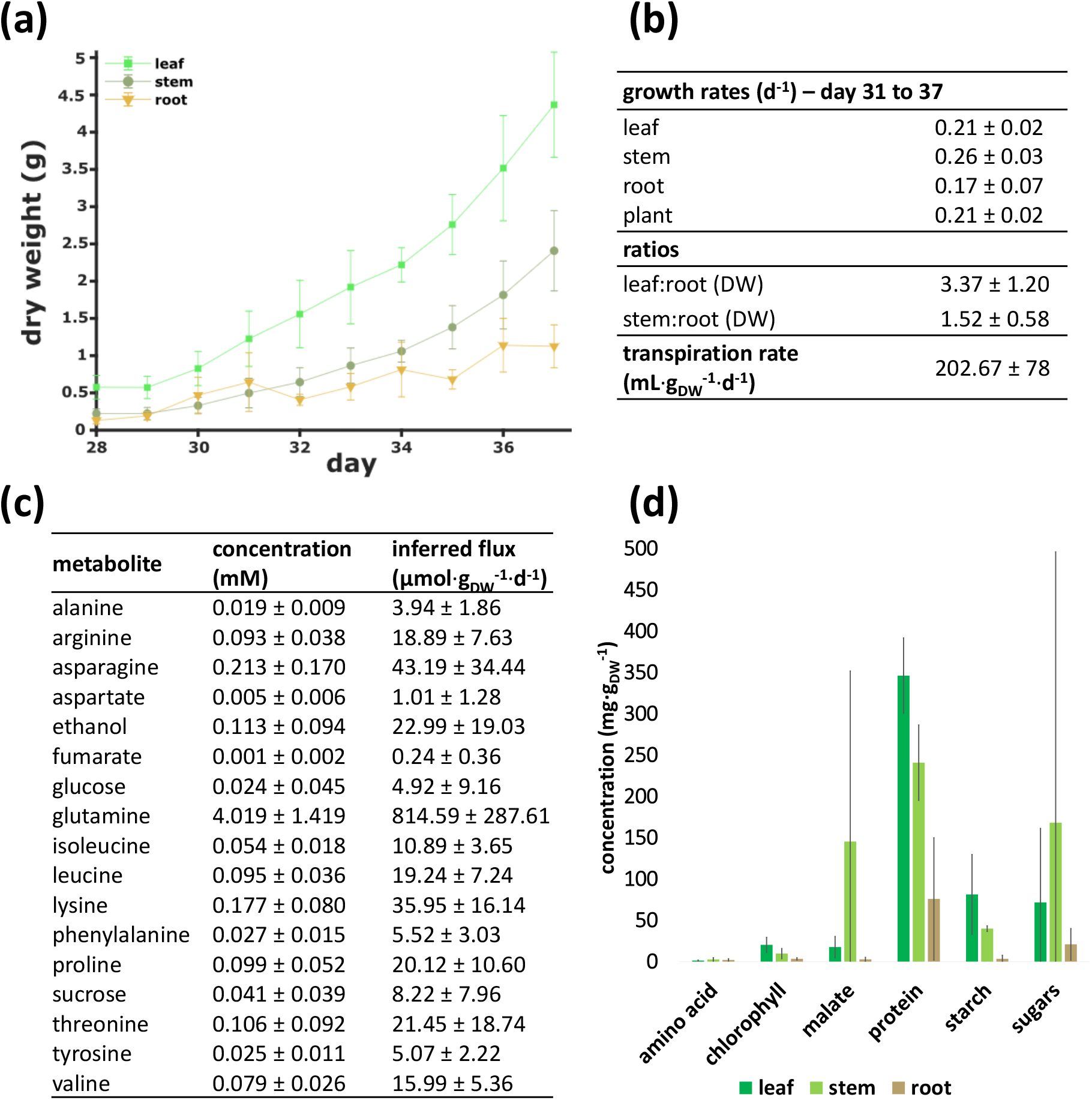
Measurement of physiological and biochemical parameters on 4-weeks tomato plants for calibration of VYTOP. a. Organ dry weight evolution between day 31 to day 37. b. Growth rate of each organ, dry weight ratios between organs, and transpiration rate. c. Average xylem sap composition. Concentrations (mean ± standard deviation) are based on 45 samples analyzed using ^1^H NMR. **d. Organ composition.** Measurements are based on at least 19 samples per organ. Mean weights are plotted and bars indicate standard deviation

Weight dataset of sampled plants was used to measure the growth rate for each organ, the organ dry weight ratios (leaf:root, and stem:root) (Figure 2a, Figure 2b), and the fresh:dry weight ratio for each organ (Supplementary Figure 1). The growth was considered as exponential between day 31 and 37 after sewing (Supplementary Figure 1) and regressions of fresh/dry, leaf/root, stem/root weights showed high correlation coefficients (Supplementary Figure 1). Growth rates were used in VYTOP to constrain biomass fluxes; organ ratios were implemented in exchange reactions to appropriately represent the difference of weight between the three organs (**Figure *1***). Transpiration was constant per mass unit, and was converted into a transpiration rate (Figure 2b). Transpiration rate was further used to compute xylem metabolite fluxes (see Materials and Methods).

We measured organ biomass composition on fresh samples of growing tomato (Figure 2d). As expected, leaf is the organ accumulating the highest proportion of starch and has the highest chlorophyll content. The data obtained was completed with literature data for lipids, minerals, (hemi)cellulose and lignin to generate accurate organ specific biomass equations (Supplementary File 1). For each organ, the biomass equation yields 1 g of dry weight biomass (Supplementary File 1).

In comparison with Yuan et al. model iHY3410 (2016), our updated biomass equation for leaf had a C:N ratio closer to the one observed from experimental data (6.5) in high CO_2_ level, high N provided as a NO_3_^−^:NH_4_^+^ mixture (Royer, Larbat, Le Bot, Adamowicz, & Robin, 2013): 5.48 in VYTOP vs 18.52 in iHY3410.

Quantification of metabolites concentration in xylem sap (Figure 2) was performed using NMR. Glutamine was the major metabolite in xylem sap (4.019 ± 1.419 mM). Fifteen other organic molecules were detected: 11 amino acids, 2 sugars, ethanol and fumarate, whose concentrations were much lower than that of glutamine (<0.250 mM). Metabolic fluxes in xylem were computed using these concentration data and the transpiration rate of plants (see Materials and Methods).

We also gathered data from literature to choose which compounds is present in the phloem compartment (Hijaz & Killiny, 2014) and to estimate photosynthetic activity in stem. Former experimental results indicated that photosynthesis in stem is limited due to less exposed surface to light (Hetherington, Smillie, & Davies, 1998). According to this study, the leaf:stem ratio of contribution to photosynthesis is around 7.7 (considering here the petioles as stems), and this value was added in the model as a constraint. Th Ammonium:nitrate uptake ratio can be optionally constrained in the model to take into account different fertilizer conditions.Ssince the fertilizer used for the experiments contained both ammonium and nitrate and we did not measure the uptake rate of each nitrogen source, the ratio was let unconstrained for our simulations except for use case 4.

### Model framework of VYTOP is validated by experimental data

We used Flux Balance Analysis (FBA) (Orth, Thiele, & Palsson, 2010), a constraint-based modeling approach to compute the metabolic fluxes in VYTOP. FBA requires a quasi-steady-state approximation (QSSA) in the whole system. This approximation is commonly accepted for bacterial growth modeling, but to extend the approach to plant vegetative growth, we checked if QSSA was still reasonable. We examined:

i. the daily weight ratios between organs and observed no significant discrepancies between days (Supplementary Figure 1),
ii. the organic chemical composition of xylem, and also observed no important daily variation (Supplementary Figure 2),
iii. the three organ growths, which followed, as expected, an exponential growth (Supplementary Figure 1),
iv. the biomass composition of each organs, and found that it had no significant daily variation (Supplementary Figure 3).

These experimental observations performed over a 10-day period proved that we could reasonably extend QSSA to a tomato plant at vegetative growth stage. We did not take into account the day/night variation as we measured plant growth on the scale of the day and not of the hour, considering that our simulation results represented an overall average of the metabolic fluxes over a 24-hour period (Bénard et al., 2015).

Simulation of metabolic fluxes using FBA implies defining a linear optimization problem: the formulation of linear mathematical constraints (equalities or inequalities) and a linear objective function (maximization or minimization of one or several fluxes). The choice of this objective function is still debated for plants (Collakova, Yen, & Senger, 2012; Sweetlove & George Ratcliffe, 2011). We assumed that plant metabolism is regulated to favor the most efficient use of light uptake and carbon fixation to grow, which was translated into minimization of the photon uptake flux, while growth was constrained via the biomass flux value. In addition, we assumed that plant metabolism was also efficient in their use of enzymes and solved a second optimization problem with as objective function minimization of the sum of all the model’s fluxes (absolute value). This second simulation was also performed to avoid stoichiometric balanced cycles (SBCs). The resulting fluxes of this double optimization problem were analyzed in the following sections.

### Use case 1: VYTOP predicts metabolic fluxes in the whole plant

Figure 3 depicts the central metabolic fluxes obtained through modeling (Supplementary File 3). We used Flux Variability Analysis (FVA) on the central metabolic fluxes to estimate how much they were constrained and if many alternative pathways existed. FVA provides for each reaction the minimum and maximum flux values that still sustain the optimal solution found in FBA (Mahadevan & Schilling, 2003). We found no variation on the fluxes obtained, except for some reactions of the glycolysis (pyruvate kinase, phosphofructokinase) that can either be performed in the chloroplast or in the cytosol (Supplementary File 3).

**Figure 3:**
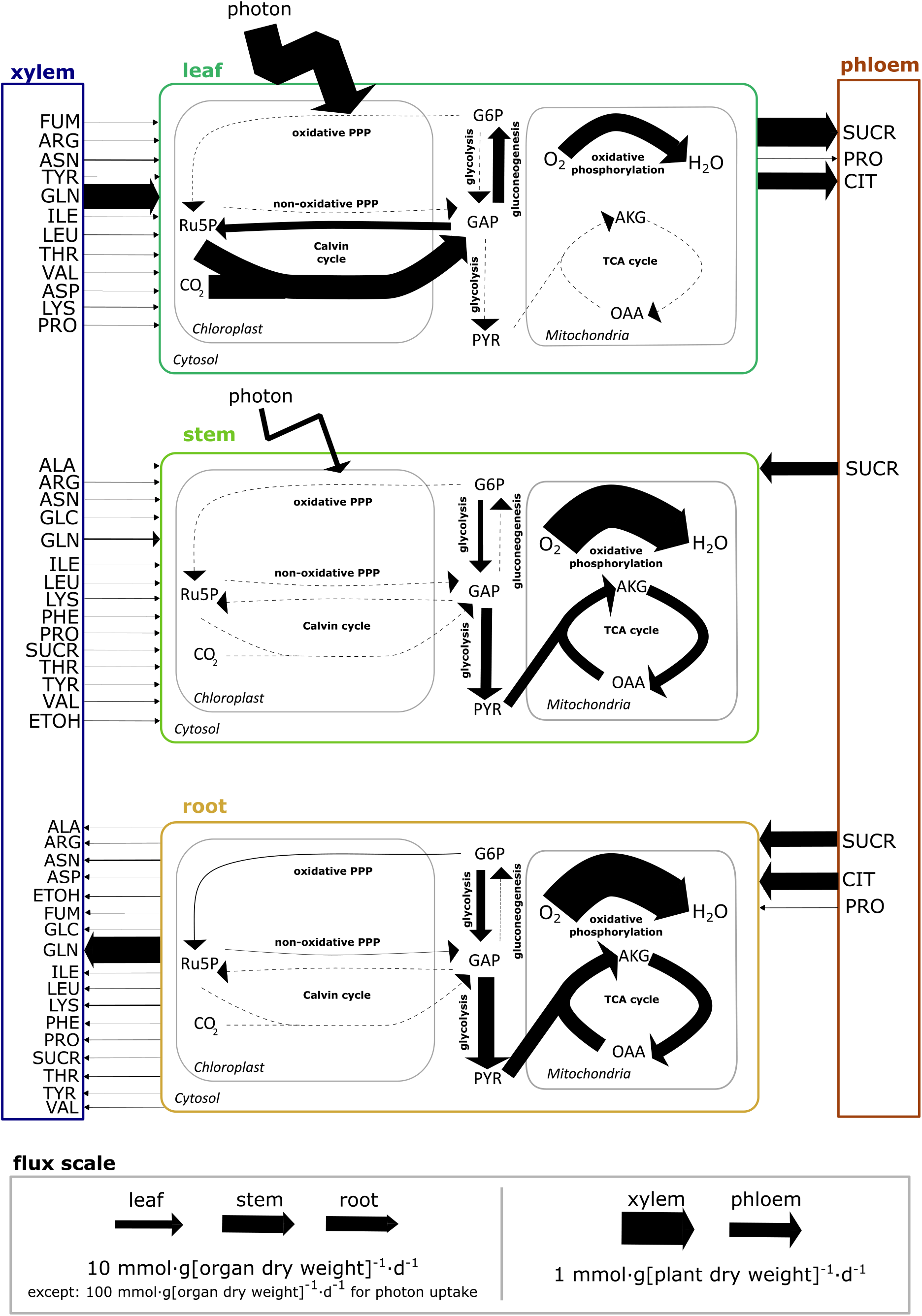
Core carbon metabolic fluxes of a tomato plant at vegetative growth stage predicted by VYTOP. Each pathway flux was estimated based on a reaction flux value representative of the pathway (list in Supplementary File 3). AKG: alpha-ketoglutarate, OAA: oxaloacetate, PYR: pyruvate, GAP: glyceraldehyde 3-phosphate, G6P: glucose-6-phosphate, Ru5P: ribulose-5-phosphate, ETOH: ethanol, SUCR: sucrose, GABA: gamma-amino-butyric acid, GLC: glucose, FUM: fumarate. Other abbreviations follow standard abbreviations for amino acids

The analyses established that core metabolic fluxes are consistent with the current knowledge on plant physiology:

i. Metabolic activity is more intense in leaves than in stems and roots. This makes sense since leaves generate the major part of organic matter necessary for the whole plant growth. In particular, there is intense photosynthetic activity and organic matter production in leaves with major fluxes in the Calvin cycle, while this pathway is not used in root and poorly used in stem.
ii. The generated organic matter is majorly converted into sugars (sucrose, glucose, fructose), either transformed into starch or transferred to stem and root through the phloem flux. Most of the organic matter (>50% of leaf assimilated carbon) is used directly by the leaf, as it is the organ with the highest weight (Figure 2).
iii. Stem needs a lower sucrose uptake flux compared to roots (40% of available sucrose versus 60%), because it has a small photosynthetic activity allowing the production of part of the needed carbon and energy (as ATP and NADPH). Yet, stem is still dependent on leaf carbon production.
iv. Glycolysis and TCA hold high fluxes in stem and root, allowing the generation of energy and biomass precursors from phloem sucrose and citrate.

Previous multi-organ metabolic models of plants were developed (Shaw & Cheung, 2020). The term multi-organ model refers here to models that represent several plant organs with different roles for each organ and with metabolite exchanges between each organ and not to tissue-specific models which simulate one type of organ at a time. For the multi-organ metabolic models developed so far, the calibration of constraints (organ ratio, growth rate, authorized fluxes) to accurately predict the behavior of a plant organ remains rudimentary and do not rely on extensive biochemical and physiological data. A question that also arises for these models are how to represent accurately metabolism differences between each organ. To this end, several frameworks were developed and rely on different strategies. Some of these frameworks such as the model of *Arabidopsis* developed by Mintz-Oron et al. (2012) used -omics data (RNAseq, proteomics, metabolomics) to take into account organ/tissue specificities. The initial methodology consisted in pruning inappropriate reactions using an expression threshold on RNAseq data to decide whether or not the reaction takes place in the organ/tissue, and therefore should tend to a closer biological relevance. However, it is limited by the fact that the transcription does not include subsequent regulation layers (translation, post-translational modifications and enzyme activation), which can be particularly important for central carbon metabolism (de Groot et al., 2007). Also, choosing this expression threshold can be sometimes arbitrary, particularly for non-model plants. Other frameworks were developed since, which minimize these drawbacks, such as GIMME-like, iMAT-like, MBA-like methods and the RegrEx method (Robaina Estévez & Nikoloski, 2014; Scheunemann, Brady, & Nikoloski, 2018). Even if thresholds can still be used in some of these methods, pruning is no as strict as the initial method since only a minimization of the discrepancy between the model and -omics data is performed. Finally, another multi-organ plant model relies on defined constraints with a global source-sink macroscopic model representing the allocation of carbon between organs (Grafahrend-Belau et al. 2013).

In our approach, we chose not to constrain each organ using -omics data, to allow all the possible chemical reactions *a priori*. To constrain VYTOP, we chose instead to use an important set of external physiological constraints (growth rates, ratios, well-calibrated biomass, photosynthetic contributions) to impose organ-specific and sink/source metabolic behaviors in the different organs, which is closer to the methodology developed for barley. The carbon fluxes distribution in central metabolism obtained in VYTOP demonstrated that this approach, relying only on physiological constraints, allows an overall good prediction of metabolic fluxes between the different organs of a whole plant, despite the non-use of -omics data. Its disadvantage is that it relies on the acquisition of physiological (and ideally metabolic) data that are not always available (whereas numerous sets of RNAseq data are available). However, it advantageously removes some of the methodological questions required to build -omics-based metabolic models (Richelle, Joshi, & Lewis, 2019). In the future, it would be interesting to compare the predictions of VYTOP with models exploiting -omics data in order to reveal more precisely the advantages and drawbacks of each methodology.

### Use case 2: VYTOP assesses the impact of physiological changes on the plant

We used VYTOP to assess the influence of the stem on plant metabolism, namely its contribution to photosynthesis (**Error! Reference source not found.**) and the effect of the stem / whole plant ratio ().

The contribution of the stem to photosynthesis represents the ability to perform photosynthesis comparatively to leaf and depends on the surface of the stem (Hetherington et al., 1998). The model shows that this contribution strongly affects photon demand since the latter decreases rapidly while the stem photosynthesis capacity increases (**Error! Reference source not found.**). In agreement with this observation, photosystem I metabolic flux strongly decreases in leaf while it increases in stem, as it becomes more advantageous to perform photosynthesis in stem (also true for photosystem II, see Supplementary Figure 4). We also observed a proportional decrease of leaf export to phloem and stem uptake from phloem. However, there is no carbon contribution of the stem to phloem, because the range of values shown here did not allow sufficient photosynthetic activity of the stem to support both stem and root growth. Yet, a sucrose flux from stem to phloem appeared when stem contribution to photosynthesis exceeds 40% (Supplementary Text 1). Simulations also showed that increasing the stem weight proportion in the plant impacted progressively the photon demand (). In this case, the plant stem plays a role of sink for carbon sources, which progressively becomes a burden affecting the photon demand (i.e synthesis of carbon sources).

VYTOP does not integrate the architectural role of the stem in plant growth, enabling a better access to light that counterbalances its cost as sink of matter. A model integrating both plant geometry and metabolic fluxes would therefore be of great interest to study the trade-off between the architectural role and the carbon sink, but remains a challenging issue.

### Use case 3: VYTOP infers the composition of tomato xylem sap by applying physiological constraints

To evaluate the relevance of VYTOP on exchange fluxes between organs, we simulated plant metabolic fluxes with the experimentally measured xylem fluxes on the root to xylem export reactions as additional constraints in the model. We analyzed how these additional constraints impacted the global photon demand (**Figure** ***6***a). We observed a very low effect on photon demand since photon demand increased by 0.63% if a constraint is set on glutamine, the major organic molecule in concentration. Photon demand increased by 0.92%, if a constraint is set for each metabolite measured experimentally. Therefore, experimental values do not disturb much the photon demand. This emphasized that VYTOP is a consistent model of matter exchange between plant organs.

**Figure 4:**
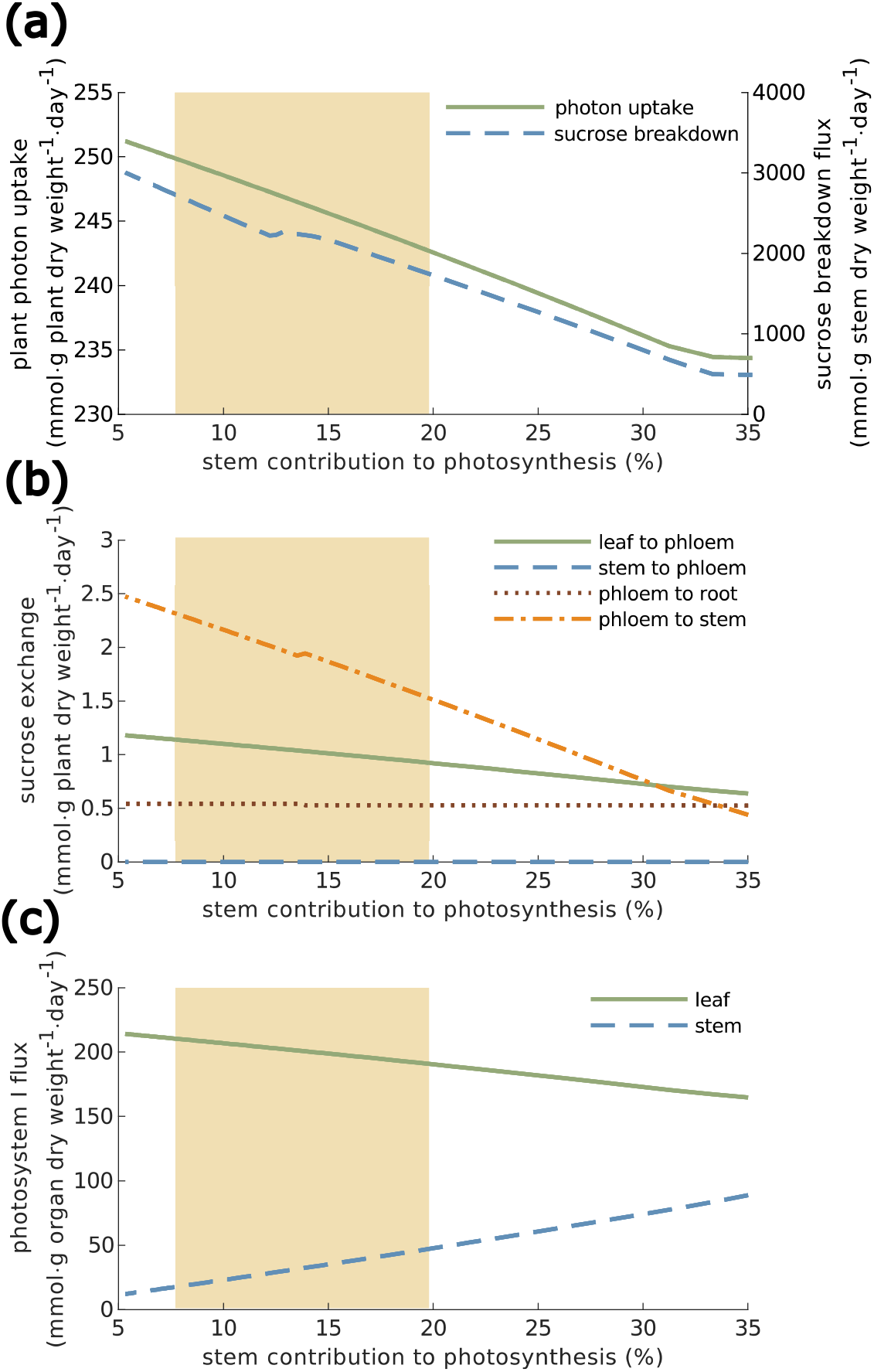
Effect of different leaf:stem photosynthetic capabilities on photon uptake (a), sucrose exchange (b) and photosystem I (c). Results are presented in percentage of stem contribution to photosynthesis. (a) Effect of stem contribution to photosynthesis on plant photon uptake and sucrose breakdown fluxes. (b) Effect of stem contribution to photosynthesis on sucrose exchange reactions between organs. (c) Effect of stem contribution to photosynthesis on the photosystem I flux. The yellow area represents the range of experimentally observed values: mean ± standard deviation.

**Figure 5.**
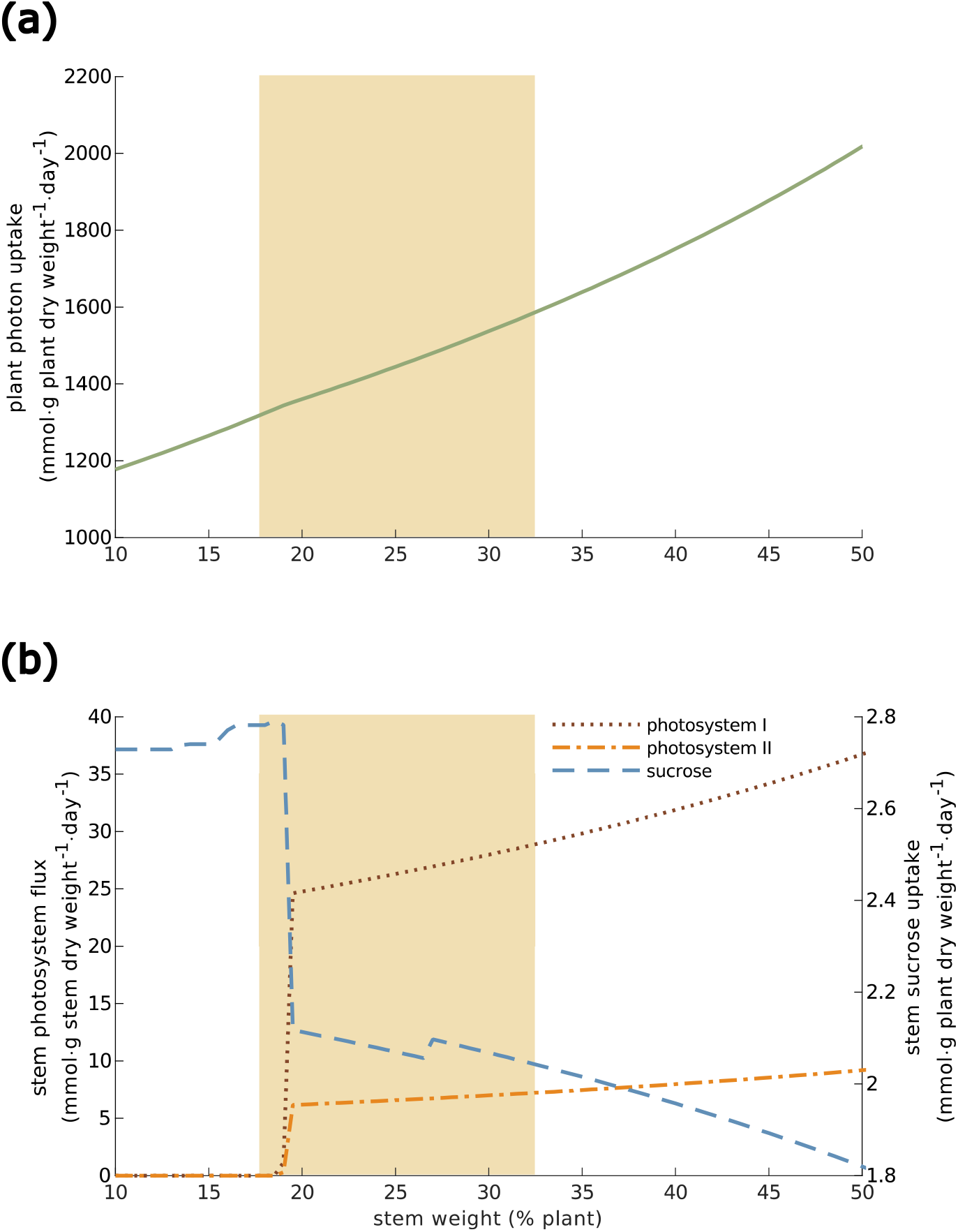
Effect of stem weight on photon uptake (a), sucrose uptake flux in stem (b) and photosystems fluxes in stem (b). The yellow area represents the range of experimentally observed values: mean ± standard deviation.

**Figure 6.**
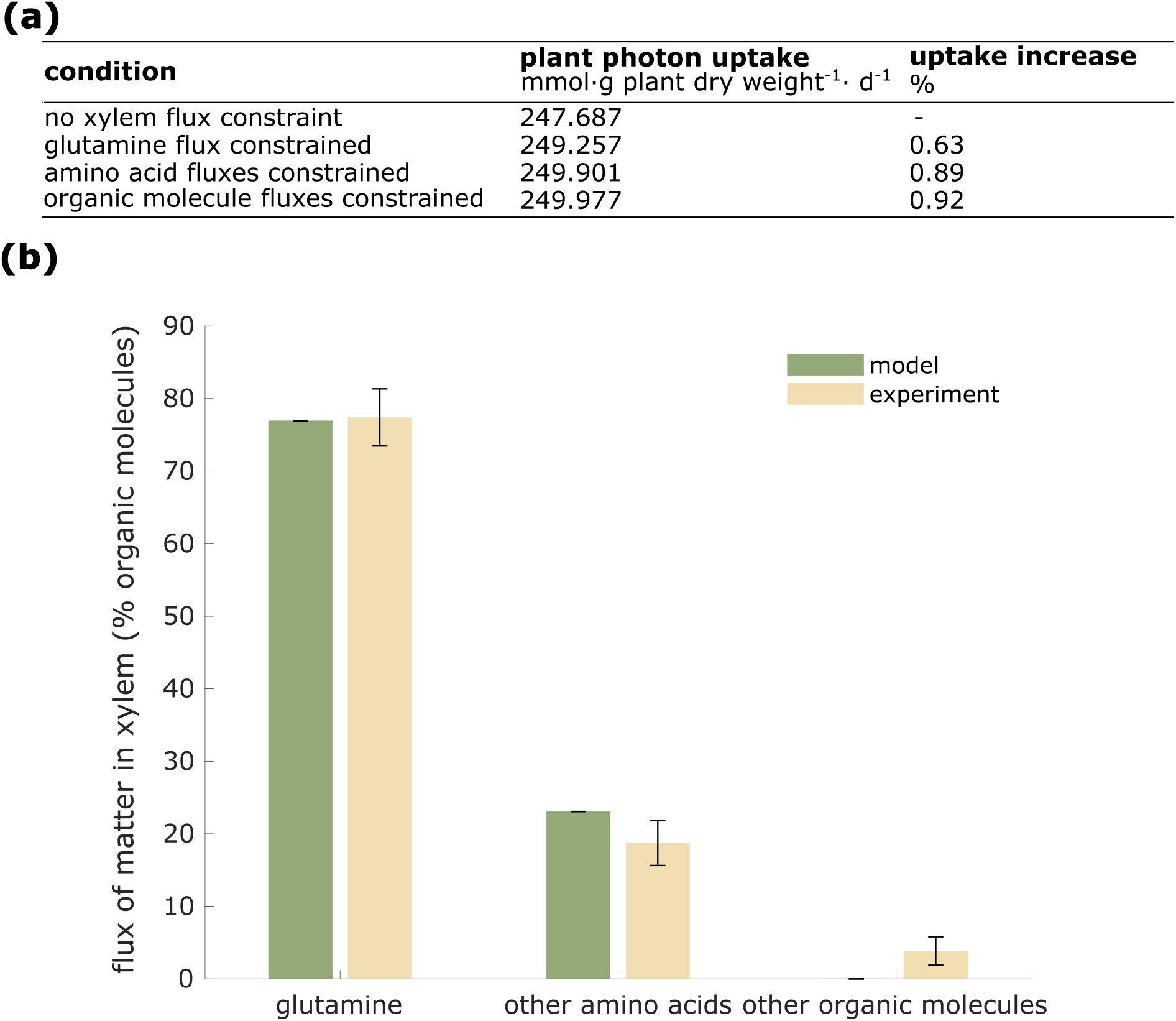
Prediction of tomato xylem fluxes by VYTOP. a. Impact of the addition of constraints on xylem fluxes (set from experimental data) on plant photon demand b. Comparison of predicted fluxes by VYTOP and experimental fluxes of organic matter in xylem sap. Modelling results were obtained from a simulation without xylem flux constraint. Error bars indicate FVA range for the modelled distribution and mean ± standard deviation range for the experimental distribution.

In a second analysis, we compared the VYTOP composition of xylem fluxes to the experimental data (**Figure** ***6***b) with, this time, no constraints on root to xylem exports. The global features of xylem chemistry are well predicted by VYTOP: predominance (around 80%) of glutamine, a central metabolite found in xylem sap from diverse plant species is inferred by VYTOP. VYTOP also predicts the presence of other amino acids, in accordance with reported experimental data (Andersen & Brodbeck, 1989; Montes Borrego, Miguel ; Jiménez-Díaz, Rafael M. ; Trapero Casas, José Luis ; Navas Cortés, Juan Antonio ; Haro, Carmen; Rivas, J. C.; Fuente, L. de la; Landa, 2017; Zuluaga, Puigvert, & Valls, 2013). FVA results (Supplementary File 3) shows no variability for the different organic fluxes predicted, as represented in **Figure** ***6***b by the absence of error bars. FVA thus confirmed that VYTOP predicts the importance of glutamine in xylem sap, as observed by metabolomics experiments.

With only global physiological constraints, specific biomass equation and resource use optimization (photon minimization and sum of flux minimization) (Figure 1, Figure 2), VYTOP efficiently predicts transport of metabolites to aerial parts. This result showhs that the composition of xylem sap is probably driven by plant physiology and resource optimization. The predominance of glutamine in both the model and experiments is remarkable. Glutamine is, with glutamate, the precursor of other amino acids and can be directly used as a carbon source since it is directly connected to the TCA cycle. Transporting a central metabolite, branched with both central metabolism and amino acid biosynthesis, thus appears more advantageous and robust. The predominance of glutamine over glutamate in xylem sap may be due to the acidic property of the latter, which could be deleterious at high concentrations. The presence of other amino acids is also a conserved property of xylem sap. We hypothesize that this redundancy of nitrogen sources could bring robustness to plant metabolism.

Several other organic molecules (organic acids, sugars, ethanol) were observed experimentally but not predicted by the presented flux distribution (Figure 6b). This must be due to a certain level of exchange and porosity between compartments that exists in nature and is not taken into account in the model, such as possible exchange fluxes between xylem and phloem via stem cells. Furthermore, an ascending phloem is observed in tomato (Bonnemain, 1980), contributing to the observation of sugars in xylem sap. Presence of ethanol may reflect an oxygen limitation in certain tomato cells, a local phenomenon not depicted in the model. Organic molecules can also act as shuttle for some ions, such as citrate for iron (Rellán-Álvarez et al., 2010). VYTOP does not represent these complex behaviors and provides a simplified and more schematized view of matter fluxes, nevertheless consistent for around 95% of xylem sap content.

### Use case 4: VYTOP predicts the impact of nutritional limitation on tomato growth rate

We used VYTOP to predict how a whole tomato plant responds to changes in a nutrient resource. De Groot et al. (2002) analyzed how the tomato growth rate was affected by nitrogen limitation. They monitored growth rates associated to different N supplies (Supplementary File 3), which revealed a progressive decrease of plant growth rate (Figure 7). To simulate the impact of N limitation in VYTOP, we used the N uptake flux and photon uptake computed in optimal conditions as constraints and progressively decreased the N uptake flux as in De Groot et al. (2002), upon the same photon availability. The simulation was performed with only nitrate as nitrogen source, as in the experimental study. We assumed as new objective function the maximization of leaf biomass growth, with constant root/leaf and stem/leaf ratio, obtained from our experimental results in nitrogen replete conditions. The simulation predicted a linear decrease of growth rate, which followed the same pattern as for the experiment, with very close values. For example, both VYTOP and the experimental data showed a reduction of the plant growth rate from 0.24 to around 0.13 day^−1^ when the nitrogen content is reduced by 50%. Thus, VYTOP agrees with the experimentally observed impact of N limitation on tomato plant growth.

**Figure 7:**
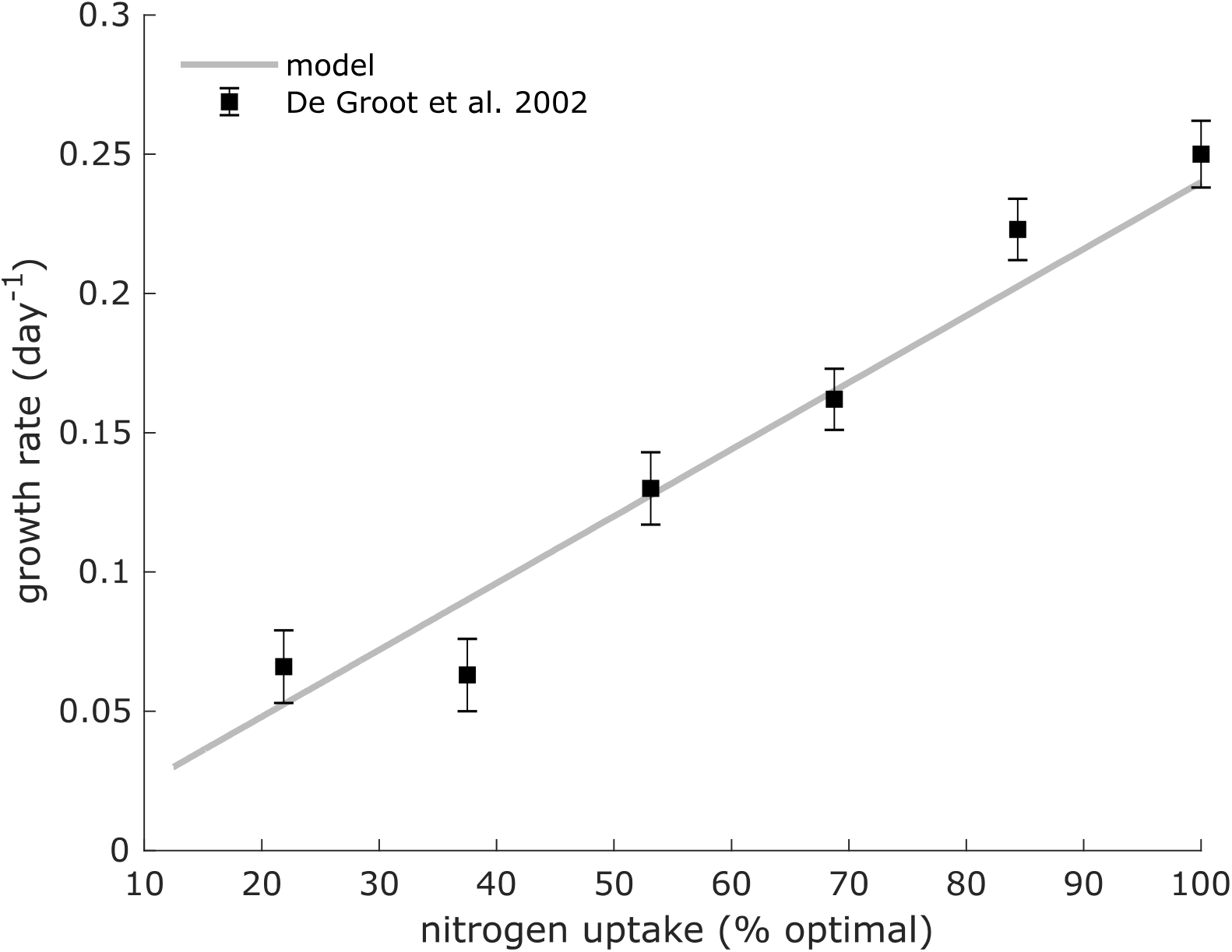
Impact of nitrogen limitation on plant growth. Experimental data were taken from De Groot et al. (2002). Growth rates were predicted using VYTOP model with a limited N uptake flux.

### Use case 5: VYTOP predicts and helps to understand the physiology and metabolism of transgenic tomato lines

We assessed how efficiently VYTOP can predict the effect of a genetic perturbation on the plant physiology (i.e mutant behavior). We selected four studies presenting transgenic tomato lines. Three lines express a fragment of a TCA cycle gene in the antisense orientation, resulting in a reduced enzymatic activity of respectively mitochondrial citrate synthase (mCS) (Sienkiewicz-Porzucek et al., 2008), mitochondrial succinyl-CoA ligase (SCoAL) (Studart-Guimarães et al., 2007) and mitochondrial alpha-ketoglutarate dehydrogrenase (AKGDH) (Araújo et al., 2012). The fourth line is a Sucrose Phosphate Synthase (SPS) gene knockout (C.M Rick TGRC https://tgrc.ucdavis.edu/Data/Acc/GenRepeater.aspx?Gene=Sus). We modeled these transgenic lines with VYTOP by forcing the matter flux to zero for the knockout line or performing simulation at 50% of the optimal flux for attenuated lines, in agreement with the activity reduction presented in the three studies. We then analyzed the impact of these changes on metabolic fluxes.

One major experimental result on the three lines with attenuated activity on TCA is that the impact on vegetative growth and young tomato physiology appears to be weak, with slight reduced growth and photosynthetic activity similar to the wild-type line (Araújo et al., 2012; Sienkiewicz-Porzucek et al., 2008; Studart-Guimarães et al., 2007). Similarly, using VYTOP, we found that plant growth was the same. We also analyzed photosynthesis flux predicted in the model and found only a very limited impact (<1.5% of flux increase) compared to the wild-type model.

In the different experimental studies, authors suggested that there was a bypass to compensate the enzyme deficiencies in the transgenic plants. In line with this hypothesis, VYTOP predicted an activation of peroxisomal citrate synthase flux for the line attenuated in mCS. The experimental study on the same line confirms this prediction since an enhanced activity of peroxisomal Citrate Synthase is observed. Similarly, for the lines attenuated in mSCoAL and mAKGDH activities, VYTOP predicts the activation of three reactions: GABA-Transaminase, Succinic Semialdehyde Dehydrogenase and Glutamate Decarboxylase, constituting the GABA shunt. This pathway can generate succinate from alpha-ketoglutarate without the use of AKGDH and SCoAL. Interestingly, metabolomics samplings and ^14^CO_2_ in the experimental studies also point toward the use of this pathway.

Conversely, no bypass exists after SPS gene knockout, as its silencing led to no growth in the resulting homozygous line. VYTOP also predicts this dependence as no growth can be obtained with silenced SPS flux. Thus, VYTOP seems to be an adequate tool to predict the alternative pathways that could be used to bypass a bottleneck in a transgenic line, or to predict the essentiality of some other reactions and the absence of alternative pathways.

VYTOP could thus be both a predictive tool to anticipate the effect of genetic engineering before generating a transgenic line, or a way to understand and analyze a phenotype observed on the transgenic line. However, it is worth noting that metabolism is often tightly linked with regulation, and that some effects are not predictable by VYTOP yet. As an example, the model could not predict reduction of nitrate metabolism observed in plants with attenuated mCS activity, and the authors suggested that it was due to posttranslational regulation. The development of hybrid metabolic-regulation models is challenging, particularly because regulation is difficult to predict and reconstruct at a genome-scale level.

### Conclusion

Modeling the complex interactions between nutritional sources, genes, energetic requirements and exchange of matter is an ongoing challenge in plant systems biology. In this study, we developed a modeling framework that integrates all these components, based on genome-scale metabolic model and physiological characterizations. The resulting tomato model VYTOP predicts consistent flux distribution and global behavior for each organ without constraining the internal fluxes, highlighting that the methodology used is relevant to build organ-specific models. We demonstrated that VYTOP gives consistent results on five different use cases: prediction of core metabolic fluxes and exchanges, role of stem in the plant, analysis of xylem composition, estimation of nutrient limitation impact on growth and impact of genetic modifications on metabolism and physiology. This tool should be useful to guide genetic/metabolic engineering of plants as well as to understand at a system level how plants respond to nutritional variations. Other modeling approaches conducted on tomato linked plant architecture and physiology to environmental variations such as temperature (Cieslak et al., 2016; Pradal, Dufour-Kowalski, Boudon, Fournier, & Godin, 2008; Pradal, Fournier, Valduriez, & Cohen-Boulakia, 2015). It will be interesting to combine these macroscopic models with VYTOP. In addition, VYTOP could be extended to other tomato plant lifecycle stages, by adding metabolic models for flower and/or fruit. Adding other metabolic models for organism interacting with plant, such as fungi or bacteria (Peyraud et al., 2017), is also a promising perspective to explore plant interactions with other organisms.

## Materials and Methods

### Experimental procedures

#### Plant cultures and automatic phenotyping

Tomato seeds (*Solanum lycopersicum* M82) were grown in soil (SB2, Proveen, The Netherlands) supplemented with Osmocote® coated fertilizer at a rate of 4g/L. Osmocote® coated fertilizer contained nitric nitrogen (5.3%), ammoniacal nitrogen (6.7%), 7% of phosphorus pentoxide, 19% of potassium oxide.

Seed were germinated in a growth chamber (26°C, 67% HR, 12h-LED light per day). Around a hundred of plantlets were transplanted in individual plastic pots (8×8cm) 8 days after sowing. 16 days after, 90 young plants were chosen and repotted in 3L pots until the end of the experiment. Foam cover discs were placed on each pot to limit physical evaporation. The plants were loaded on the Phenoserre robot facility of the Toulouse Plant Microbe Phenotyping infrastructure. 12h-light per day at 28°C and 50% humidity were programmed.

All the plants were watered with 100 ml three times on the loading day and weighted, in order to define a well-watered target weight. The daily transpiration was determined as the weight differences between two consecutive days, at the time of watering. Transpiration rate was deduced as following: transpiration rate at day i (mL·g[dry weight]^−1^·day^−1^) = transpiration at day i (mL·day^−1^) / plant dry weight at day i (g[dry weight]). Temperature, hygrometry and light intensity were recorded during the whole experiment.

#### Collection and preparation of plant samples

Six plants were removed each day from the conveyor belt for samplings during 9 consecutive days, starting 4 days after the loading on the robot. For these plants, stems were cut just above the cotyledons node, rinsed with approximately 1 ml of water and the upcoming xylem sap was harvested by repeated pipetting and collection into Eppendorf tubes placed on ice. The tubes were placed at −80°C for further quantitative NMR analyses.

#### Quantitative NMR analyses

The xylem saps were analyzed by 1D 1H NMR on MetaToul analytics platform (UMR5504, UMR792, CNRS, INRAE, INSA 135 Avenue de Rangueil 31077 Toulouse Cedex 04, France), using the Bruker Avance 800 MHz equipped with an ATMA 5mm cryoprobe. Each xylem sap sample was centrifuged to remove the residues (5min, 13520 RCF, Hettich Mikro 200 centrifuge), then placed in 3 mm NMR tubes. TSP-d4 standard (Sodium 3-(trimethylsilyl)(1-13C,2H4)propanoate) was used as a reference. pH 6.0 phosphate buffer was used to standardize the chemical shifts among samples. Acquisition conditions were as follows: 30° pulse angle, 20.0287 ppm spectral width, 64 scans per acquisition for a total scan time of approximately 8 minutes per sample, and zgpr30 water pre-saturation sequence. The samples were kept at a temperature of 280 K (6.85 °C) all along the analysis. Resonances of metabolites were manually integrated and the concentrations were calculated based on the number of equivalent protons for each integrated signal and on the TSP final concentration.

Xylem sampling and NMR analysis allows only to have access to the concentration of a metabolite (in mmol·mL-1) at a given time. However, xylem sap is a continuous flow rate. Using the transpiration of the plant, and assuming a constant concentration of the metabolite in the xylem on a 24h basis (validated throughout our experiment, see section “Model framework of VYTOP is validated by experimental data”), the flow of metabolite per day can be estimated using the following formulae:

Metabolite flux (in mmol·g dry weight^−1^·day^−1^) = Metabolite concentration (in mmol·mL^−1^)· transpiration rate (mL·g dry weight^−1^.day^−1^).

#### Biochemical analyses of metabolites

The different organs of each plant were collected separately (stems, leaves, roots) on another experiment performed on the same platform and plant variety and close plant age (4-week old). Approximately 300 mg fresh weight for each collected organ was frozen in liquid nitrogen and stored at −20°C for further biochemical analyses.

Quantifications of metabolites were performed at the HiTMe platform (INRAE - IBVM - 71 avenue E. Bourlaux - CS 20032 - 33882 Villenave d’Ornon Cedex, France). The plants samples, previously frozen in liquid nitrogen, were ground to a powder using liquid nitrogen to avoid thawing. A quantity of 20 ± 10 mg of each were weighted in previously frozen Micronic tubes. Free amino acids, glucose, fructose, malate, proteins, starch, sucrose and chlorophylls in leaves, stems and roots were quantified as described in (Biais et al., 2014). Briefly, ethanolic extracts from every samples were obtained using three consecutive incubation of the frozen ground powder aliquots. Ethanol 80% v/v with HEPES/KOH 10 mM pH6 buffer was used for the two first incubations, and ethanol 50% v/v with HEPES/KOH 10 mM pH6 buffer was used for the third. Supernatants were pooled and used for the quantification of chlorophylls, glucose, fructose, sucrose, malate and free amino acids. Pellets were used for the determination of protein and starch contents. The extracts and pellets were stored at −20°C between each quantification. For each sample, chlorophylls were quantified by measuring optical densities at 645 and 665 nm on a mix of 50 μl of extract supplemented with 150 μl of analytics grade ethanol. Amino acids were quantified using the fluorescamine method. Excitation wavelength was 405nm and emission was measured at 485 nm. The proteins were quantified using Bradford reagent. Starch was quantified in glucose equivalent after full pellet digestion in an oven at 37°C for 18 hours. For the other analytes cited above, the NADH/NADPH appearance was measured, and the analytes were quantified using a 1:1 stoichiometric coefficient.

### *In silico* procedures

#### Tomato metabolic network curation and conversion

The tomato metabolic model iHY3410 published by Yuan et al. (2016) was used as starting point. The authors provided tomato metabolic network in two formats: table (.xlsx) and SBML (Hucka et al., 2003). Some reaction directions were different in the SBML and table files, and some metabolites (proton and water) were missing in the table file. Thus, we performed a first curation step to merge the files and select the appropriate reaction directions.

Then, we converted the metabolic network identifiers (IDs) (metabolites and reactions) to BiGG nomenclature. The advantage of BiGG nomenclature is that it is usually more explicit for the analysis as the IDs are usually an abbreviation of the metabolite/reaction names, while in MetaCyc ID it is often a generic acronym (CPD for metabolites, RXN for reactions) and a number. The web tool SAMIR (Semi Automatic Metabolic Identifier Reconciliation) (Peyraud, Cottret, Marmiesse, Gouzy, & Genin, 2016) was used to find the appropriate BiGG IDs for metabolites and reactions. The correspondences between MetaCyc and BiGG IDs are provided in the final metabolic network in table format (Supplementary File 1). For some metabolites and reactions, no BiGG ID was found even after manual researches in BiGG database, probably because they are specific to plants. New BiGG IDs following the usual nomenclature were created for 1094 reactions and 758 metabolites (see Supplementary File 1). The SBML file (Hucka et al., 2003) was generated using Met4j (https://forgemia.inra.fr/metexplore/met4j) and ModelPolisher (Römer et al., 2016).

The resulting metabolic model was tested on autotrophic growth and heterotrophic growth (using FBA, see Simulations section) on different organic substrates before generating a multi-organ metabolic model. Growth was achievable with CO_2_ + light, starch, sucrose, glucose, fructose, glycerol, glutamate, glutamine, alpha-ketoglutarate, fumarate, asparagine, alanine, isocitrate as carbon sources. Other organic (glutamine, glutamate, aspartate, asparagine, alanine, proline, histidine, leucine, methionine, lysine, cysteine) and inorganic (nitrate, ammonium) elements can be used as nitrogen sources. No catabolic reactions were found for the assimilation of lysine, ethanol, isoleucine, beta-alanine, leucine, cysteine as carbon sources whereas they were experimentally observed in the xylem in amount superior to biomass assimilation, suggesting that they could be catabolized. Thus, we searched the catabolic pathways of these amino acids in plants and incorporated them in the network, using BLAST to find the orthologous proteins in *S. lycopersicum*. 21 reactions, linked to 34 genes were added to the model iHY3410. 15 reactions had a GPR complete with an e-value superior to 5e-131 (Supplementary File 1), 6 had none. After this second curation step, we computed the mass balance (on carbon, nitrogen, phosphate, sulfur and oxygen) and manually curated the reactions with incorrect carbon, nitrogen, phosphate, sulfur and oxygen balance. We generated a final tomato metabolic network Sl2183. The model is available in MetExplore (Cottret et al., 2018) https://metexplore.toulouse.inrae.fr/metexplore2/?idBioSource=6237, BioModels (Glont et al., 2018) under the ID xxx and in a github repository https://github.com/lgerlin/slyc-metabolic-model/ [once the paper is accepted].

#### VYTOP construction

To generate VYTOP (Virtual Young TOmato Plant) (see flow chart in Supplementary Figure 5), we built organ-specific metabolic models for leaf, stem and root. To this end, organ growth rates and organ-specific biomass equations were determined from our experiments and implemented as constraints in the simulations. We generated an in-house script to parse the metabolic network Sl2183 (SBML file) and replicate it in three to represent three organs. In the generated model, each organ is represented as a “metacompartment”: metabolite and reaction IDs has a final letter (l s and r for leaf stem and root respectively) to indicate their organ location (e.g R_NAD2 reaction becomes R_NAD2_l, R_NAD2_s, R_NAD2_r and M_gln_L_c becomes M_gln_L_c_l, M_gln_L_c_s, M_gln_L_c_r).

Our in-house script also generates exchange reactions to represent transfers between organs and exchange compartments (represented with the letters xyl for xylem and phl for phloem), such as 1 M_gln_L_c_r -> *n* M_gln_L_xyl and 1 M_gln_L_xyl -> *m* M_gln_L_c_l. The list of metabolites authorized to be exchanged in xylem and phloem respectively is available in Supplementary File 1. *n* and *m* represent the mass ratios between the organs and the whole plant, in order to take into account the different weights of organs and have quantitative predictions at the whole plant level. For example, for transfer from the root to the xylem, 1 μmol/g root dry weight/day will generate *n* = 1/(1 + 1.52 + 3.37) = 0.1697 μmol/g plant dry weight/day in xylem, and then 1 μmol/g plant dry weight/day in xylem will generate m = (1 + 1.52 + 3.37)/3.37 = 1.74 μmol/g leaf dry weight/day. Finally, a transport cost (ATP cost) of 0.5 was added in the model: transporting k μmol/g organ dry weight/day would consume k·(transport cost) μmol/g organ dry weigh/day of ATP. Performing several simulations of use case 3 with different value of ATP transport cost, we found that a transport cost between 0.02 and 1.0 mol of ATP per mol of exchanged molecule, which appears reasonable, provided the most consistent predictions in agreement with experimental data, while use case 4 is not impacted by the different set of values tested (Supplementary Text 1).

Physiological constraints were added to impose different physiological roles in organs.

i. Uptake of minerals and water is not allowed in leaf and stem, while it is allowed with no limitation in roots (assuming they are not limiting factors for well-watered plant and adequate nutritional supply in the soil).
ii. Photon uptake was allowed in leaf and stem but not in root.
iii. Ammonium:nitrate uptake ratio can be constrained. Few data are available to predict the balance between ammonium and nitrate in root uptake. We analyzed the impact of varying the parameter in Supplementary Text 1. The ratio was not constrained in our simulations, excepted use case 4, to agree with the experimental data
iv. Photosynthesis is limited in stems by its reduced exchange surface compared to leaves (Hetherington et al., 1998). We then introduced a ratio (leaf/stem) of contribution to photosynthesis, to limit the ability of stem to perform photosynthesis: photosynthesis in stem is constrained to be inferior or equal to photosynthesis in leaves / surface ratio.
v. Photorespiration was modelled by imposing a ratio between rubisco carboxylase and oxygenase fluxes. The ratio was set to 85, which represents the mean ratio of enzymatic specificity for CO_2_ versus O_2_ for C3 plants, according to experimental data gathered by Tcherkez et al. (2006).

Finally, ATP maintenance, which represents the global cost in a cell for non-metabolic processes (such as housekeeping functions), was kept at the value used by Yuan et al. (2016) as no organ specific ATP maintenance values are available.

#### Computational simulations

Modeling of a whole tomato plant metabolism was performed using constrained-based modeling, with the Flux Balance Analysis methodology (Orth et al., 2010). Briefly, FBA relies on i) the quasi-steady-state assumption (QSSA) in the biological system modeled ii) the formulation of a biologically relevant objective: flux(es) to minimize/maximize iii) eventually additional physiological constraints. The solution of FBA will be the optimal matter fluxes (regarding the objective) in a metabolic network assuming QSSA and respecting the constraints. Use of QSSA in a whole tomato plant is justified by our experimental results (see part “Model framework of VYTOP is validated by experimental data” of the Results & Discussion section). We decided to define the minimization of total photon uptake as our objective. The photon uptake flux obtained from this simulation was then integrated as a new constraint in the model, and FBA was run a second time with the objective of minimizing the sum of absolute fluxes. The use of these two objectives is discussed in the Results and Discussion section part ‘Model framework of VYTOP is validated by experimental data’.

FBA was run with or without the integration of experimentally measured xylem fluxes, and with varying stem proportions, surface ratios and transport costs.

Flux Variability Analysis (FVA) (Mahadevan & Schilling, 2003) was also performed. This extension of the FBA aims at finding alternative solutions of the one generated by FBA, as its flux distribution is usually not unique. It consists in imposing the optimal objective values found in FBA (here both photon uptake and sum of absolute fluxes) as additional constraints, and determining the minimum and maximum flux that can carry each reaction in these conditions.

Simulations were performed with Python 3.5 scripts, the open access libraries lxml, pandas and the linear programming solver CPLEX Python API, developed by IBM and free for academic institutions. Scripts are available at https://github.com/lgerlin/slyc-metabolic-model [once the paper is accepted].

**Table 1:** Effects of transgenic tomato lines on plant physiology and metabolism and comparison between experimentally observed effects and VYTOP predictions. mCS: mitochondrial citrase synthase, SCoAL: succinyl-CoA ligase, AKGDH: alpha-ketoglutarate dehydrogenase, SPS: sucrose phosphate synthase, pCS: peroxisomal citrate synthase. Experimental results for mCS, SCoAL, AKGDH and SPS are respectively from Sienkiewicz-Porzucek et al. (2008), Studart-Guimarães et al., (2007), Araújo et al. (2012) and C.M Rick TGRC (https://tgrc.ucdavis.edu/Data/Acc/GenRepeater.aspx?Gene=Sus).

| Enzyme | Activity | Effect | VYTOP prediction | Consistency |
| --- | --- | --- | --- | --- |
| mCS | Attenuated | Minor effect on growth | Optimal growth achieved | + |
| mCS | Attenuated | Unaltered leaf photosynthesis | <1.5% increase | + |
| mCS | Attenuated | Enhanced CSp activity | CSp activation | + |
| mCS | Attenuated | Reduction of nitrate metabolism | No change | - |
| SCoAL | Attenuated | Minor effect on growth | Optimal growth achieved | + |
| SCoAL | Attenuated | Unaltered leaf photosynthesis | <1.5% increase | + |
| SCoAL | Attenuated | Evidences of GABA shunt use | GABA shunt activation | + |
| AKGDH | Attenuated | Unaltered leaf photosynthesis | <1.5% flux increase | + |
| AKGDH | Attenuated | Evidences of GABA shunt use | GABA shunt activation | + |
| AKGDH | Attenuated | Minor effect on growth | Optimal growth achieved | + |
| SPS | Suppressed | Lethal | Growth infeasible | + |

## Supporting Information

**Supplementary Figure 1. Additional physiological data.** Fresh:dry weight ratio for each organ, growth curves with ln(dry weight), linear regression of fresh/dry weights and organ weights, daily variation of transpiration.

**Supplementary Figure 2. Additional xylem metabolomics data.** Xylem constitution at each sampling day.

**Supplementary Figure 3. Additional organ metabolomics data.** Biomass constitution at each sampling day.

**Supplementary Figure 4. Additional representation of model responses to plant physical variations.** Metabolic fluxes in photosystem II as a function of stem contribution to photosynthesis.

**Supplementary Figure 5. Flow chart of the multi-organ modeling pipeline.**

**Supplementary File 1. Table file of tomato plant genome-scale metabolic network Sl2183.** Contains gene-protein-reaction (GPR) information, BiGG/MetaCyc associations, biomass equation details for each tissue, lists of reactions and metabolites, new BiGG IDs generated for the model.

**Supplementary File 2. SBML file of tomato plant genome-scale metabolic network Sl2183.** SBML Level 3 version 1.

**Supplementary File 3. Table file with modeling results.** List of reactions used to represent carbon core metabolic fluxes and FVA on central metabolism reactions. De Groot et al (2002) and Sienkiewicz-Porzucek et al (2008) data used for model validations. List of reactions used to predict mCS reduced activity impacts.

**Supplementary Text 1. Additional analyses.** Effects of ATP cost value, nitrogen uptake ratio and physiological changes at high percentages, analysis of SBCs.

## Acknowledgments

We thank TPMP (Toulouse Plant Microbe Phenotyping) platform (Castanet-Tolosan, France) and its staff Nemo Peeters, Felicià Maviane Macia and Fabrice Devoilles for their technical support in plant cultures and imaging, as well as HiTMe (High-Throughput Metabolic phenotyping) platform (Villenave d’Ornon, France) for biochemical analysis of organ metabolites. We acknowledge MetaToul (Metabolomics & Fluxomics Facitilies, Toulouse, France, www.metatoul.fr) platform, which is part of the MetaboHUB-ANR-11-INBS-0010 national infrastructure (www.metabohub.fr) and its staff Cécilia Berges, Edern Cahoreau and Lindsay Peyriga for access to NMR facilities. We also thank Sophie Colombié for helpful discussions and feedbacks on the manuscript.

## Data Availability

All data supporting the findings of this study are available within the paper and within its supplementary materials published online.

